# Combined Atomic Force Microscope and Volumetric Light Sheet System for Mechanobiology

**DOI:** 10.1101/812396

**Authors:** E. Nelsen, C.M. Hobson, M.E. Kern, J.P. Hsiao, E.T. O’Brien, T. Watanabe, B.M. Condon, M. Boyce, S. Grinstein, K.M. Hahn, M.R. Falvo, R. Superfine

**Affiliations:** Dept. of Physics and Astronomy, University of North Carolina at Chapel Hill, Chapel Hill, NC 27701; Dept. of Pharmacology, University of North Carolina at Chapel Hill, Chapel Hill, NC 27701; Dept. of Biochemistry, Duke University School of Medicine, Durham, NC 27710, United States; Program in Cell Biology, Hospital for Sick Children, Toronto, ON, M5G 0A4, Canada; Dept. of Applied and Materials Science, University of North Carolina at Chapel Hill, Chapel Hill, NC 27701

**Author notes:** Falvo and Superfine are co-corresponding authors.

## Abstract

The central goals of mechanobiology are to understand how cells generate force and how they respond to environmental mechanical stimuli. A full picture of these processes requires high-resolution, volumetric imaging with time-correlated force measurements. Here we present an instrument that combines an open-top, single-objective light sheet fluorescence microscope with an atomic force microscope (AFM), providing simultaneous volumetric imaging with high spatiotemporal resolution and high dynamic range force capability (10 pN – 100 nN). With this system we have captured lysosome trafficking, vimentin nuclear caging, and actin dynamics on the order of one second per volume. To showcase the unique advantages of combining Line Bessel light sheet imaging with AFM, we measured the forces exerted by a macrophage during FcɣR-mediated phagocytosis while performing both sequential two-color, fixed plane and volumetric imaging of F-actin. This unique instrument allows for a myriad of novel studies investigating the coupling of cellular dynamics and mechanical forces.

## MAIN

Cells interact mechanically with their environment by generating and responding to forces. A focus on the mechanical dynamics of cell phenomena such as motility, division and phagocytosis is essential for understanding stem cell fate ^1^, cancer progression ^2^ and innate immunity ^3^. These mechanical processes are inherently three-dimensional (3D) and are regulated both by very local (nm) interactions as well as whole-cell scale (μm) biochemical and mechanical signaling. Obtaining a more complete picture of a cell’s mechanical interaction with its environment requires monitoring local and global structures in 3D while simultaneously measuring associated forces. These goals necessitate integrating high spatial and temporal resolution volumetric imaging methods with high resolution force acquisition. Light sheet florescence microscopy (LSFM) enables acquisition of volumetric, multicolor time series at a high resolution and frame rate with low background fluorescence and low phototoxicity ^4–6^. Among single cell force methods ^7^, Atomic Force Microscopy (AFM) ^8^ is unique in combining a large force range (10^−11^-10^−6^ N) that enables molecular-scale to tissue-level mechanics measurements, with high bandwidth temporal resolution (μs) and sub-nanometer spatial control. Combining LSFM with AFM (AFM-LS) imposes significant geometrical constraints to the optical system design, and demands low vibration operation to accommodate sensitive force measurements. To address this challenge, we used a single-objective selective plane microscopy (soSPIM) technique integrated with, and time-synchronized to, an AFM. As described in a report on our first generation system ^9^, we generated a fixed vertical light sheet through the objective while collecting a side-ways view (x-z) fluorescence image through the same objective with the aid of a 45 degree mirrored prism. Here we describe our second generation system that combines vertical Line-Bessel Sheet (LBS) illumination with volumetric image collection for multicolor, 3D fluorescence movies that are time-correlated with sensitive force measurements.

The second generation system comprises several advances that enable high frame rate volumetric imaging with accompanying force application and measurement. Full computer control of the optical path by electronically controlled steering mirrors and electrically tunable lenses (ETL) on the input and output sides provide high frame rate sheet scanning and image focusing capabilities. The system employs a remote focusing technique that uses an ETL on the detection path to change the focal plane without moving the objective lens. This minimized the mechanical vibrations that would otherwise compromise force measurements. To provide extended depth of field for the desired light sheet width ^10^, a LBS was implemented ^11^. We collected two-color, 4D (3D + time) volumetric scans with the LBS on live cells with simultaneous, time-synchronized AFM force data acquisition. Our single cell 3D scans were collected on the order of one second, which is comparable with reports of similar fluorescence imaging techniques ^10,12^.

To challenge our imaging system, we investigated several contexts for which high quality visualization of 4D cytoskeletal dynamics is essential to address the critical biophysical questions. To demonstrate the volumetric imaging capabilities of our system, we focused on vimentin. Vimentin is a representative component of the intermediate filament (IF) cytoskeleton, a network with viscoelastic properties ^13,14^ and functions distinct from those of the actin and microtubule systems ^15^. IFs are essential for normal cell morphology, motility and signal transduction, and IF dysregulation underlies over 80 human diseases ^14^. Despite its importance, we understand relatively little about the mechanobiology, dynamics and regulation of the IF network, as compared to other cytoskeletal components ^15,16^. Current techniques are unable to connect optical measurements of assembly state dynamics with force measurements to probe the mechanical properties of IFs in real time. Using our system, we monitored the 3D architecture of the vimentin cytoskeleton in human cells on a single-second timescale, demonstrating sufficient temporal and spatial resolution to capture IF assembly state dynamics. Additionally, we demonstrate the 4D data collection capabilities of our system by presenting data of lysosome movement within the vimentin network.

To demonstrate our combined imaging and force capabilities, we investigated the biophysics of Fcγ receptor-mediated phagocytosis by macrophages. Phagocytosis is the process by which large pathogen particles are engaged through specific ligand-receptor interactions, engulfed, and then digested ^17^. Although the details of actin structural dynamics during phagocytosis have been investigated ^18–20^, it is increasingly clear that forces and mechanosensitivity are also integral to immune cell function, a research area recently dubbed “mechanoimmunology” ^3^. We present simultaneous volumetric image and force data acquisition of actin cytoskeleton dynamics in the phagocytic cup during target engulfment. With the AFM-LS, we were able to test current hypotheses of the time course of outward and inward engulfment forces ^21^, and their relation to actin remodeling ^22^. The relative timing, spatial structure and magnitudes of the macrophage forces and cytoskeletal dynamics may be critical in understanding the connection between morphological and biochemical events that occur during the phagocytic process. To this end, we also present quantitative cross-correlation of engulfment force and local actin dynamics.

## RESULTS

### AFM-LS design

Our system combines high spatiotemporal resolution 3D fluorescence microscopy with single-molecule level force spectroscopy; LSFM provides the former while AFM is capable of the latter (Fig. 1, **Supplementary Fig. 1** and **Supplementary Fig. 2**). We have previously described implementation of single-objective LSFM in a conventional microscope ^23^. In brief, a vertical LBS was generated at the sample plane by relaying the image of an elliptically shaped beam intersecting an annular aperture to the back focal plane of the objective. A right-angle prism was lowered adjacent to a cell of interest and the image plane was raised, using the objective focus, to the height on the prism where the virtual image of the rotated view of the specimen is formed. The x-y virtual image is the x-z plane of the cell as it was illuminated by the vertical light sheet (See **Supplementary Fig. 3** for side-view imaging optical theory and **Supplementary Fig. 4** for point spread function). ETLa and ETLb (Fig. 1a, **Supplementary Fig. 1**) control axial scanning of the illumination light and image plane respectively; the scanning mirror (Fig. 1a, **Supplementary Fig. 1**) translated the light sheet at the sample plane. We can achieve either full frame rate fixed light sheet side-view imaging of an x-z plane, or sequentially offset x-z slices to produce volumetric imaging of an entire cell.

**Figure 1.**
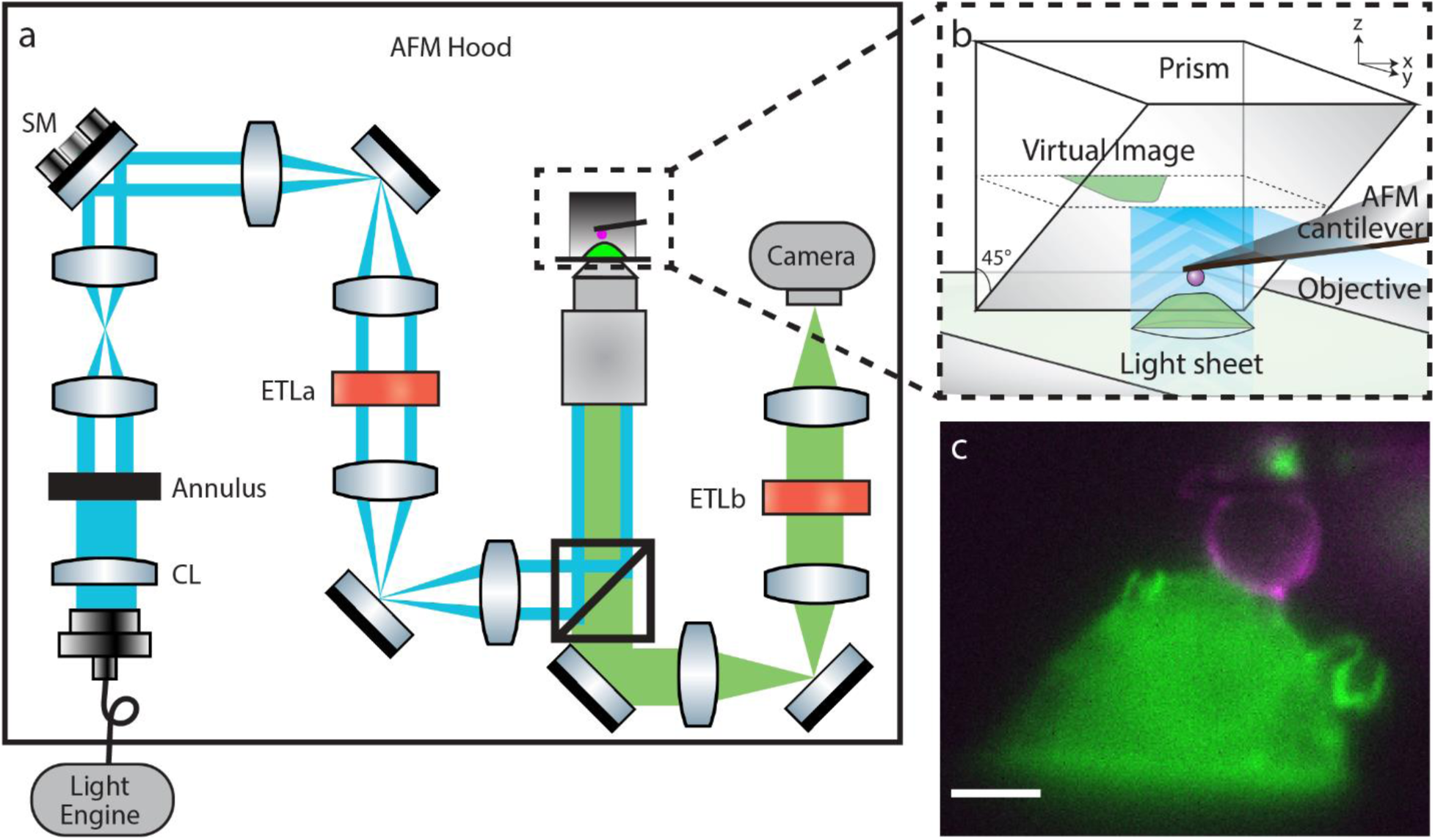
Schematic of the combined AFM and LSFM system. **a**, Laser lines housed in our light engine are coupled into an optical fiber that passes into the AFM hood. A cylindrical lens focuses a sheet of light onto an annulus which is re-imaged to the back focal plane (BFP) of the objective to generate a Line Bessel Sheet. ETLa and ETLb control the axial position of the light sheet and image plane respectively. The steering mirror (SM), conjugate to the BFP of the objective, controls the position of the light sheet at the sample plane (See also **Supplementary Figure 1**). **b**, Schematic of the integration of the prism and AFM in the sample space. **c**, Side-view LSFM image of a RAW 264.7 cell expressing HaloTag F-tractin labeled with Janelia Fluor 549 (green) and a polystyrene bead attached to an AFM cantilever coated with fluorescent IgG (magenta). Scale bar = 5 µm.

Integration of the AFM into the optical system is shown in Fig. 1b. The AFM cantilever was positioned between the cell and the prism ^9^. All of the beam-shaping optics were designed to fit within the AFM hood, and the beam scanning was tuned to have no measurable effect on the AFM force noise, which at ~10 pN, is sufficient for molecular scale force measurements (**Supplementary Fig. 5**). Each dynamic optical component was computer-controlled, which allowed us to seal the acoustic hood enclosing the system throughout the duration of each experiment (see **Supplementary Fig. 2**). Fig. 1c illustrates a typical side-view image of a cell, in this case a RAW 264.7 macrophage. The two-color image shows the fluorescence of Halo-tagged F-tractin in green and a 6 μm polystyrene bead attached to an AFM cantilever and coated with fluorescent IgG (magenta), touching the top of the cell.

### Two-color volumetric imaging

Our system is capable of performing fast (~1 s), single-objective, two-color LSFM volumetric imaging (Fig. 2, **Supplementary Video 1** and **Supplementary Video 2**). We first performed volumetric imaging of a live HeLa cell stably expressing vimentin-mEmerald and labeled with Lysotracker Deep Red (Fig. 2a, **Supplementary Video 1**). This temporal resolution allowed us to visualize the 3D dynamics of lysosomes within the vimentin network on single-second timescales. This opens up the potential to study lysosome transport dynamics and their interaction with the cytoskeleton ^24^ in cells under controlled mechanical load. Recent reports show that the vimentin cytoskeleton plays a central role in regulating lysosome trafficking in autophagy ^25^.

**Figure 2.**
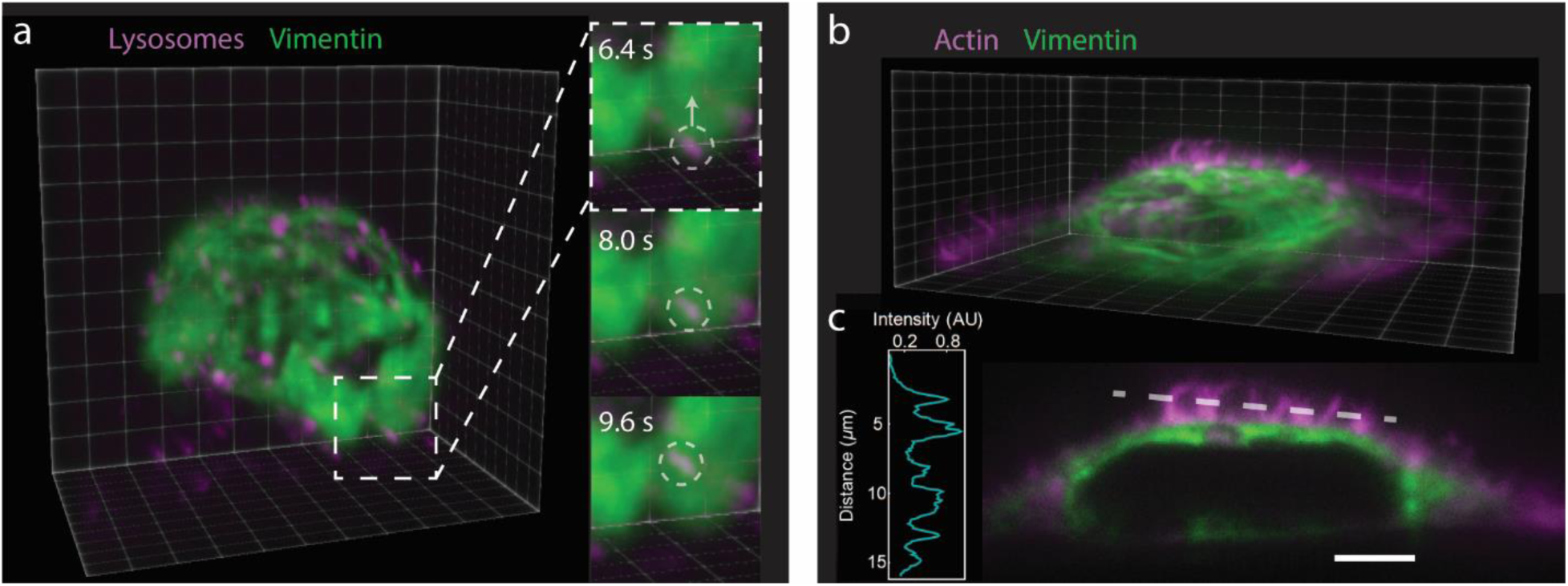
Volumetric imaging of vimentin cytoskeleton and lysosome movement. **a**, LSFM volumetric imaging of a live HeLa cell with labeled Lysotracker Red (magenta) and vimentin-mEmerald (green). Each volume consists of 75 slices per color acquired at 5 ms exposure time and 5 ms readout time; the total volume acquisition time is 1.5 s. A 100 ms delay is taken between volumes. Insets show dynamics of vesicles on the single-second timescale. Voxel size: 106 × 106 × 220 nm. Total volume: 29.3 × 26.3 × 16.5 µm (See **Supplementary Video 1**). **b**, LSFM volumetric image of a fixed HeLa cell with labeled actin (AlexaFluor 568-Phalloidin - magenta) and vimentin-mEmerald (green). The volume consists of 300 slices per color acquired at 5 ms exposure time and 5 ms readout time; the total volume acquisition time is 6 s. Voxel size: 106 × 106 × 108 nm. Total volume: 52.6 × 19.1 × 32.4 µm (See **Supplementary Video 2**). **c**, Selected side-view image from the volume data set shown in (b). A single-pixel line scan (dashed white line) through the actin channel shows we resolve individual filopodia at the above volume acquisition rate with adequate signal-to-noise (inset). Scale bar = 5 µm.

To illustrate the ability of the system to visualize fine detail while scanning at a high frame rate, we fixed the mEmerald vimentin cells and stained them with AlexaFluor 594 phalloidin for F-actin. We then performed volumetric imaging as before, but with one half the axial step size (Fig. 2b,c, **Supplementary Video 2**). A single-pixel line scan through the filopodia (Fig. 2c) demonstrates the signal-to-background and feature resolution our system can achieve at this scan rate with a reasonable laser intensity for live-cell studies (see Methods). Two-color volumetric imaging paves the way for experiments to characterize the mechanobiological role of IFs in processes such as organelle motility ^26,27^ and filopodia dynamics ^28^.

### Forces during phagocytosis

We employed the AFM-LS system to investigate forces generated during phagocytosis. An antibody-coated 6 μm diameter bead was attached to an AFM cantilever, and was lowered to ~0.5 µm above the top of the cell (Fig. 3a). The force applied by the macrophage (RAW 264.7) on the bead was measured as the cell sampled, and then subsequently attempted to engulf the bead. As in Fig. 1c, the macrophage was stably expressing F-actin marker F-tractin and the bead was coated with fluorescent IgG (AlexaFluor 488). We first describe fixed light sheet experiments (Fig. 3 and **Supplementary Videos 3 and 4**). Sequential images were recorded at 1.25 frames/s over the duration of ~30-40 min for each experiment, and synchronized with the AFM force data. We focused first on forces generated early in the phagocytic process. Forces generated by the macrophage early in the process are of interest as they may promote ligand-receptor engagement ^29^ and facilitate target stiffness sensing^3^. Second, we directly compared cup progression and force to test models of phagocytic cup generated engulfment mechanics. To make this comparison, we generated a kymograph of the structure of the cup around the bead’s circumference, as measured by F-tractin fluorescence (see Fig. 3).

**Figure 3.**
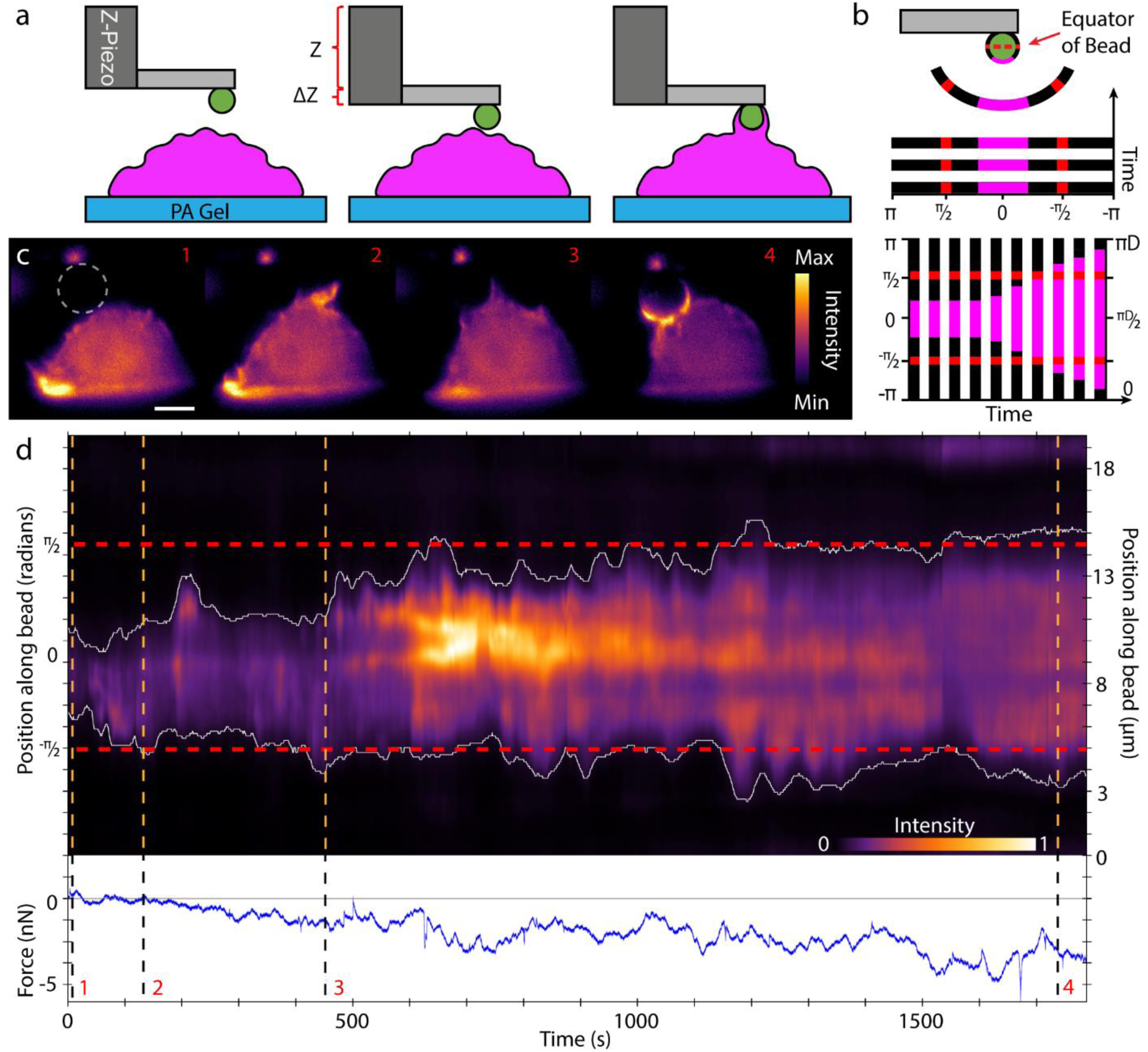
Side-view visualization and force characterization of phagocytic cup formation using LSFM and AFM. **a**, A cartoon of the experimental procedure. The base end of the cantilever is lowered using the z-piezo such that the bead is located < 1 um above the cell. Holding the z-piezo position fixed for a 30 min dwell, the cell is imaged in side view and forces are measured as the macrophage attempts to engulf the bead. We note that the bead is affixed to the much larger cantilever and therefore cannot be fully engulfed. **b**, A circular kymograph is formed by drawing a 5 pixel thick circle around the bead circumference and plotting the average intensity across this 5 pixel line vs. position along the circle. The unfurled intensity plots for each image frame are then put together in a kymograph as depicted in bottom representation of b. The red dashed line marks the equator of the bead tip. **c**, A fixed LS image sequence of the RAW 264.7 cell stably expressing HaloTag F-tractin labeled with JaneliaFluor 549 and illuminated using a 561 nm vertical line Bessel light sheet (see also **Supplementary Video 3**). Each image corresponds to its labelled orange dashed line in (d): 1, the cell before phagocytic cup formation, 2, at initial cup formation, 3, point where cup reaches the bead equator, 4, mature cup showing punctate actin. Scale bar = 5 µm. Bead tip was coated in AlexaFluor488-conjugated IgG, and was imaged in a separate channel with 488 nm laser (bead channel not shown). Each channel was imaged for 300 ms with 100 ms delay between channels. **d**, Circular kymograph depicting the actin intensity around the bead tip over the course of the experiment (retraction data not shown). Note for intensity scale for **d** is optimized for clarity and does not correspond to that of **c**. Position along circumference is represented in radians on left axis and in µm on right axis. The dashed horizontal red lines in the kymograph represent the equator of the bead. The white lines represent the edge of the phagocytic cup, taken as 11% of the maximum intensity. Lower part of the kymograph represents the right side of the bead.

Fig. 3c shows four time points of the side-view fixed light sheet experiment, with corresponding dashed lines in Fig 3d. The force on the bead vs. time is graphed below the kymograph, with the negative force direction chosen as downward (cell-ward). From the pattern of forces and extent of the cup formation, we observed in this particular experiment that significant downward force (> 1 nN in magnitude) was generated by the cell well before the cup reached the equator of the bead. In 11 of 20 experiments, we observed downward forces (> 100 pN) exerted by cells before the bead equator was reached. We also observed that in 10 of 20 experiments, an upward force greater than 50 pN was observed before the cup formed, most likely related to the macrophage probing the phagocytic target with ruffling structures or filopodia. We note for context that mechanochemical signaling at cell matrix adhesions can occur with forces below 10 pN ^30^. For the data set depicted in Fig. 3, we found a maximum of 4 nN of force which corresponds to an average downward surface stress over the bead hemisphere of approximately 70 pN/μm^2^ (70 Pa). These values are similar to mean traction stresses measured for adherent and motile cells as measured with flexible post ^31^ or traction force substrates ^32^. Other studies have shown forces within the phagocytic cup include compressive forces ^33^ on the target near the cup edge consistent with an actomyosin contractile ring ^18^. In our data, we did not find an obvious change in the direction or magnitude of the force as the cup approached and passed the midway of the target as would be expected if the contractile force at the edge dominated the net force on the target. Although the net downward force did undergo considerable fluctuations, on average the force continued to increase in the downward direction during the entire cup progression. Cell-ward forces from initial stages of cup formation were in agreement with observations and subsequent modeling performed by *Herant, et al* and *van Zon, et al*. ^34,35^.

### Correlation of force with local actin dynamics

Our side-view videos of phagocytosis (**Supplementary Videos 3 and 4**) showed that the actin cytoskeleton was dynamic across the entire cross section of the cell with significant structural features (membrane ruffles and filopodia) forming well away from the cup region. We were interested in how F-actin accumulations fluctuated during engulfment, both around the cup and in regions distal to the cup, and how this related to engulfment force. We therefore collected dynamic F-actin intensity within several distinct cell regions during phagocytosis, and cross-correlated those data with the bead engulfment force (see Online Methods for details). Because F-tractin binds only to F-actin, fluorescence intensity is taken as a proxy for local F-actin concentration. We hypothesized that F-actin concentration fluctuations within critical cell regions (e.g. the cup) were related to our measured force via combinations of protrusive polymerization forces and acto-myosin based, contractile activity. Three full engulfment experiments, performed as described for Fig. 3, were analyzed for such correlations at the cup, and in regions away from the cup, as shown in Fig. 4a, **panel 3**. We show a representative analysis in Fig. 4, for a different cell than that shown in Fig. 3. Fig. 4b shows the raw intensity data (average intensity over region) from the three regions, along with the force data.

**Figure 4.**
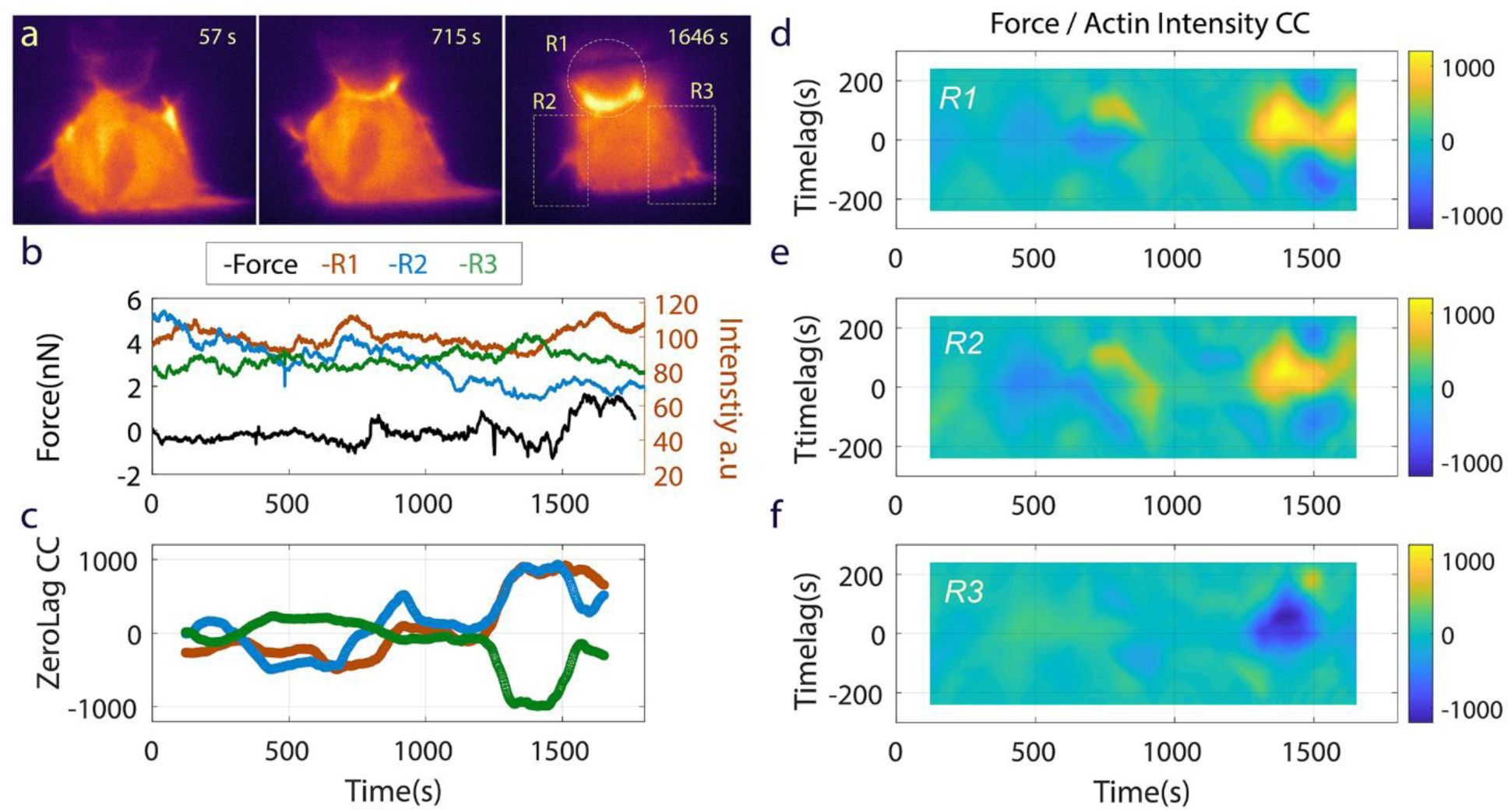
Correlation of Force and Actin Dynamics. **a**, Three regions of the macrophage were selected for actin intensity analysis: ***region 1 (R1)***, the cup, ***region 2 (R2)***, the left edge and ***region 3 (R3)**,* right edge of cell where much ruffling and filopodia activity were present (See **Supplementary Video 4**). Note the regions are non-overlapping. **b**, Raw force data and intensity data time series for the corresponding image above. **c**, Time evolution correlation coefficient for force and intensity for each region at zero time lag. The cross correlation calculation (see text for details) was done on a 240 s window of data at increments of 2 s over the whole data set. Both force and intensity data were high pass filtered prior to cross correlation calculation to remove offset and slope (frequency cut off = 0.5 mHz). **d-f**, show the correlation strength as a function of time lag across the whole duration of the phagocytic process for each region.

Note in Fig. 4c that the cup region (R1) and left-hand cell edge (R2) initially showed weaker negative correlation (at zero time lag) prior to and early in cup development (~700 s) while the right-hand cell edge (R3) was weakly positively correlated. The negative cross-correlation here at the beginning of cup development (Fig. 4a, 715 s), indicated higher actin concentration corresponding to a downward force. As the cup matured and actin intensity spiked in R1 (especially after 1500 s), regions R1 and R2 showed strong positive correlation between force and actin intensity (upward force with high actin concentration) while region R3 showed the reverse. The cross-correlation analysis also allowed us to assess relative timing of local F-actin concentration fluctuations and fluctuation in the total force on the bead. Fig. 4d-f depicts the cross-correlation data for each region as a 2D plot with the y-axis as time lag and the x-axis as lab time. Note similarity in patterns for data from R1 and R2 and the inverse pattern in R3. The fainter blue signal (corresponding to negative correlation) on the zero time lag axis at ~700 s for R1 indicates downward force coincident with initial cup formation, which is consistent with data in Fig 4c. We observed this particular correlation in all three cells for which this analysis was performed – early cup formation as indicated by increased F-actin density in the cup region coincided with downward force. Focusing next on the F-actin spike in the cup that occurred after 1500 s, we observed that the actin concentration (R1, Fig. 4d) preceded the upward force by approximately 60 s. This was similar to the correlation in region R2 and to the timing of the actin concentration of R3, although R3 displayed a negative correlation. This was also generally consistent across all three data sets: large actin intensity events in the region of the cup were strongly correlated with forces on the bead. However, later in the process after early cup formation, the direction of the forces, upward or downward, and the correlations of the local actin concentrations were variable (sometime positively correlative, sometime negatively). These results suggest that in the period following early cup formation, there is complex cell-wide coordination of actin dynamics that give rise to competing force mechanisms. This is consistent with the contributions from both actin polymerization forces and actomyosin contractile forces. For example, we hypothesize that myosin II activity is more highly correlated in time and location with downward-directed engulfment forces and less correlated with upward-directed protrusive forces. Future studies using fluorescent imaging tools to highlight myosin activity and actin polymerization will be the next step in delineating how each force generating mechanism contributes.

### Volumetric Imaging with Forces during Phagocytosis

We employed volumetric imaging to obtain a more complete view of actin structure during phagocytosis. We collected a series of volume scans of a macrophage engaging and attempting to engulf a bead. RAW 264.7 cells were prepared and labeled as described for Fig. 3. Fig. 5 presents selected volume frames and the corresponding forces during engulfment. The image sequence in Fig. 5a depicts the process of target engagement and cup formation. Fig. 5b shows the force data associated with the volumetric sequence with red lines that correspond to the frames shown in Fig. 5a. We note that here, as in the experiments of Fig. 3 and Fig. 4, there was an initial downward force of approximately 1 nN soon after macrophage engagement. Volume imaging provided a view of the F-actin signal throughout the volume of the cell. As seen in Fig. 5c and **Supplementary Videos 5 & 6**, an x-y plane maximum intensity projection of the image data in the volume of the cup showed actin accumulations, spaced ~1.5 µm from each other, at the base of the bead. These podosome-like structures ^29^ are consistent with observations reported by Barger et al ^36^ in their study investigating the interplay between myosin-I and actin structure in the phagocytic cup. In the future we will perform 3D correlation analysis to understand the role of these structures in force generation on target cells.

**Figure 5.**
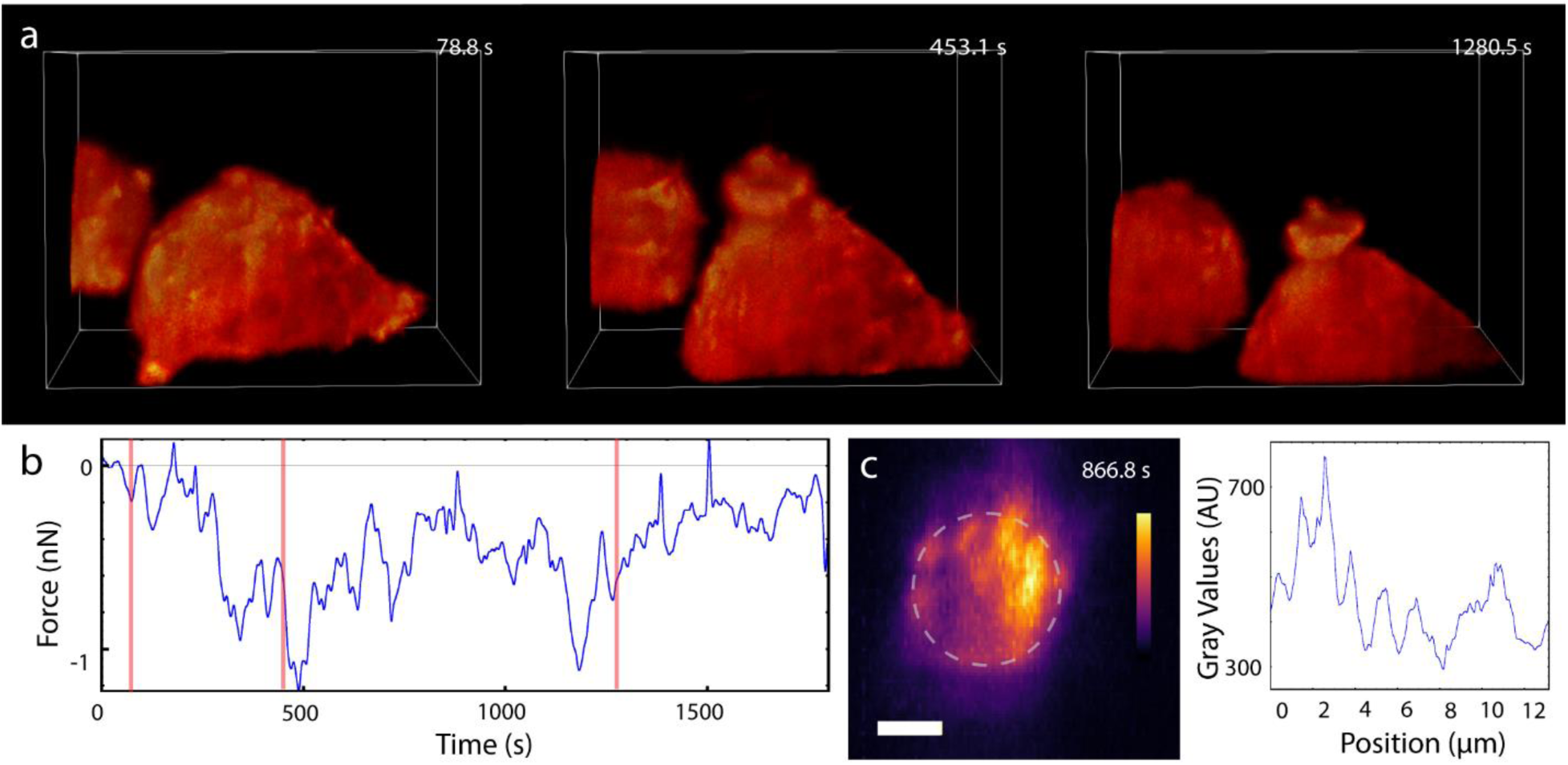
LSFM volumetric imaging and force measurement of macrophage phagocytosis. **a**, Image sequence of actin dynamics during phagocytosis. RAW 264.7 cells expressing HaloTag F-tractin labelled with JaneliaFluor 549 and the beaded tip, labelled with AlexaFluor488-conjugated IgG, were illuminated with a 561 nm and 488 nm vertical line Bessel light sheet, respectively (see also **Supplementary Video 5**). Voxel size: 106 × 106 × 200 nm. Total volume: 28.1 × 21.7 × 19.0 µm. Time stamps represent the time at the end of the volume scan. Each slice was imaged for 10 ms with 5 ms allotted readout time. Each two-color volume is 2.85 s in total with 7 s delay between colors to reduce photobleaching and phototoxicity for the half hour experiment duration. The first image corresponds to a point before the cup has started, the second to a point where the cup is forming and protruding up the bead, and the third, to a point where the cup has matured. Parts of a second cell show up in the background. **b**, Force data synchronized to the imaging. A lowpass filter with cut off frequency of 0.064 Hz was applied to the data for visualization purposes. Red lines on the force curve mark the corresponding volume frames in (a). **c**, A maximal image projection in the xy plane with accompanying intensity plot captured at 866.8 s shows bright, punctate actin structures are regularly separated by ~1.5 µm around and underneath the bead tip. Scale bar = 2 µm. (see also **Supplementary Video 6**).

## DISCUSSION

We have combined a vertical light sheet, single objective system that provides fast side-view (top to bottom) volumetric imaging simultaneous with AFM for mechanobiology applications. The implementation of a reflecting prism within the field of view provides good light collection and convenient specimen access for force probes. The LBS allows high aspect ratio illumination for low levels of phototoxicity, which enables long-term imaging. Through the complete computer control of microscope operation and careful amelioration of instrumental vibrations, we achieved single-molecule force sensitivity during image acquisition (**Supplementary Fig. 5**). The primary disadvantage of our system is the loss of light collection efficiency when imaging in side-view. The signal to noise ratio is reduced by approximately a factor of two in comparison to plan-view due to the reduced cone of light intercepted by the reflecting prism, a recognized difficulty in high viewing-angle, single objective imaging systems. The resolution suffers as well, depending upon how far the cell is from the prism. We find this is an acceptable trade-off given our systems other advantages. These issues can be fixed by employing horizontal light sheet illumination by reflecting the light sheet off the prism, with the imaging focus of the objective at the plane of the horizontal light sheet ^23^.

We demonstrated volumetric imaging of vimentin and actin networks (Fig. 2b,c) and presented 4D data revealing the movement of lysosomes within the IF cytoskeleton (Fig. 2a). We anticipate that the ability to couple force measurements with high-resolution, volumetric optical imaging will be a powerful tool for studying the mechanobiology and regulation of the IF cytoskeleton, which remain poorly understood ^27,37^. For example, vimentin filaments organize around the nucleus (Fig. 2b,c) and are known to stiffen in response to deforming forces ^38^, but the mechanism of stiffening is not yet known ^15^. Our system, combined with known vimentin mutants that limit assembly state ^39^, will allow us to interrogate this question directly. Our results also set the stage for new discoveries on IF cytoskeleton regulation. All IF proteins are decorated with post-translational modifications (PTMs), including phosphorylation and O-glycosylation, which may alter IF function ^37,40^. We previously demonstrated that the site-specific O-glycosylation of vimentin is required for IF assembly and function in human cells ^37^, but the real-time effects of these PTMs on IF network dynamics and mechanical properties remain elusive. Using simultaneous AFM-LS, we are poised to characterize the 4D effects of chemical or genetic inhibition of PTMs (e.g., phosphorylation, O-glycosylation) on the vimentin cytoskeleton, addressing a longstanding question in the IF field. In the case of lysosomes, our 4D data along with the force application capabilities we have demonstrated opens up the potential to study how load-induced distortions of cytoskeletal networks influence vesicle movement ^24^ as well as other cellular transport dynamics.

Our application of AFM-LS to the biophysics of phagocytosis revealed complexity in engulfment related actin dynamics, and in force progression. During the initial macrophage engagement, the macrophage introduced downward forces in excess of 100 pN, of sufficient magnitude for mechanoreceptor induced signaling ^30^. For cases where the cup progressed, these downward forces exceeded 4 nN in some trials. Our actin dynamics data showed strong location dependent correlation with engulfment forces, and implicated not only the actin dynamics in the cup region, but also regions distal from the cup. The correlation analysis also revealed complexity in the interplay of actin concentrations, and suggests that coordinated cell-wide cytoskeletal dynamics result in the forces imposed on the target. This in turn suggests mechanotransductive signaling during phagocytosis may be happening not only within the cup itself, but also across the cell at its basal contact, and potentially within the nucleus, with downstream consequences for cell motility, adhesion, and gene expression ^41^.

## ONLINE METHODS

### Optics for LSFM setup

The light sheet fluorescent microscope (LSFM) comprises five optical modules; four modules are in the illumination pathway and one in the imaging pathway. The optical set up with parts list and components can be found in our previous publication ^23^. We implemented modules 1-5 for the work reported here. **Supplementary Fig. 1** and **Supplementary Fig. 2** provide detailed schematic and pictures of the system. Illumination lasers (561 nm and 488 nm, OBIS, Coherent) are combined colinearly into an acousto-optic tunable filter (AOTF) (AOTnC-400.650-TN, AA optoelectronic, France) to control intensity and precise timing of each laser. A fiber optic cable (FT030-Y, Thorlabs) guides laser light into the AFM hood where the beam is shaped and relayed to the specimen through the four modules. In Module 1, the beam is expanded using beam expanders. In Module 2, the beam is focused using a cylindrical lens (LJ1567RM-A, Thorlabs) into an annulus (R1DF200, Thorlabs) forming the Line Bessel sheet (LBS). In Module 3, the lateral steering mirror (OIM 101 1”, Optics-in-Motion) controls the motion of the LBS at the specimen, enabling volume scans. In Module 4, a 4f system comprising two relay lenses and ETLa (EL-16-40-TC, Optotune) in the middle controls the axial motion of the LBS independently of the objective height. Then the beam passes through a tube lens and the objective lens to the specimen. In the imaging pathway, Module 5, a second 4f system with ETLb (EL-16-40-TC, Optotune) in the middle controls the image plane independently of the objective height. Images were captured using a Hamamatsu Orca Flash 4.0 V2+ sCMOS. A dual band pass filter cube assembly was used to pass 488 nm and 561 nm laser light to the sample and GFP and RFP emission to the camera (TRF59904-EMOL2 C173202; Chroma, USA). A short pass filter was used in the imaging path to remove light from the AFM super luminescent diode (880nm). A UplanSAPO 60x/1.2 NA W (Olympus, USA) with PIV controlled heater (HT10k; TC200; Thorlabs, USA) was used for imaging.

### Production of polyacrylamide gels

To eliminate glare from the coverslip surface that interfered with side-view imaging, we plated cells on a ~10 µm thick pad of 55 kPa polyacrylamide (PA). Bottom coverslips (40 mm round, #1; Fisher Scientific) were cleaned in a UV cleaner and vapor treated with 0.1% APTES (aminopropyl triethoxysilane; Sigma) in toluene at 80 deg for 30 min. Top coverslips (22 × 22 mm #1.5; Fisher, USA) were silanized with HMDS (hexamethyldisilane) for 30 min at 80 deg. PA gel solution was prepared with standard reagents, but with the addition of 1% polyacrylic acid v/v to provide carboxylic acid groups for linkage of ECM proteins ^33^ Quickly, 2 µL APS was added to 125 µL of gel solution, vortexed for 2 seconds, and 10 µL were placed on the center of each round, 40 mm coverslip. A 22 × 22 mm coverslip was then placed on top of the PA to spread it evenly, and a 3 g weight was placed in the center to compress it. Deionized water was placed at the edges of the 22 mm coverslip to keep the gel hydrated and aid in release. A corner of the coverslip was slowly lifted to separate the square coverslip from the gel, leaving the gel pad on the round coverslip. The round coverslip with gel pad was then placed in a biosafety hood for 15 minutes and allowed to dry enough so that a 10 mm cloning cylinder could be affixed to the gel pad with a small coating of vacuum grease (High Vacuum grease, Dow Corning, USA). An EDAC solution (1-Ethyl-3-(3-dimethylaminopropyl)carbodiimide; 10 mg/mL) in PBS was then added to the cloning cylinders for 15 min at 37 deg. The PBS/EDAC was then removed, and 10 ug/mL fibronectin in PBS was added for 60 min at 37°C. This was replaced by PBS for storage, or media for immediate use with cells.

### Prism Mounting and Cleaning

Right angle prisms (POC part 8531-607-1) were mounted onto a shaved glass capillary tube secured to a Thorlabs compact table clamp (Thorlabs CL3). A thin layer of UV polymerizing glue (Norland optical adhesive 81) was used to glue the bottom square side to the capillary tube, exposing the mirrored surface. The prism and capillary tube were then cured for 5 minutes.

Prisms were rinsed with 70% ethanol and DI water after each use to remove any accumulated debris. If further cleaning was necessary, Red First Contact polymer cleaner (Photonic Cleaning, WI) was placed onto the surface of the prism, allowed to cure for 5 minutes, then peeled off with tweezers. Repeat cleanings were performed if needed.

### Prism alignment

A right-angle prism was secured to a thin capillary tube and mounted to the prism holder stage equipped with a tip, tilt, and rotation stage (TTR001, Thorlabs). The pitch, roll, and yaw adjustments to the prism were made before each experiment using a coverslip with fluorescent beads to align the light sheet with the prism. Using translation micrometers, the prism was positioned over but not touching the AFM cantilever and cell. The relative position of the prism and light sheet was adjusted such that ETLa had enough working range to position the waist of the light sheet in the specimen while imaging in side-view. All pitch, roll, and yaw adjustments to the prism were done with the prism holder stage.

### Bead functionalization and bead attachment to AFM cantilever

6.1 µm carboxylate beads (Polysciences, Inc, USA) were dried on a cleaned coverslip coated in tetra-chlorosilane (Gilest, USA). Using the AFM head manual z translation stage (Asylum Research MFP3D, Oxford Instruments, UK), a tipless cantilever (Nanoworld TL1-Arrow-50; Nanoworld, Switzerland) was lowered onto a droplet of UV polymerizing glue (Norland optical adhesive 81; Norland Products, USA) on the same coverslip. The cantilever was lowered onto a suitable bead and a UV flashlight was used to slightly cure the glue before raising the beaded cantilever off the surface of the coverslip. The entire chip was placed under a UV lamp for 5 minutes to cure the glue completely.

The beaded cantilever was then placed in the chip holder and onto the AFM head. The tip was then dipped into a series of four droplets on a UV-cleaned coverslip: (1) a droplet of 10 mg/mL EDAC (1-ethyl-3-(−3-dimethylaminopropyl) carbodimide; Fisher Scientific) in PBS for 2 min; (2) PBS, 2 min; (3) 33 ug/mL Goat anti-Mouse IgG-AlexaFluor488 (ab175473, Abcam) for 5 min; (4) PBS, >2 min and submerged until used. IgG was centrifuged for 5 min at 10,000g before use in functionalization steps.

### Cantilever Calibration

Calibration of the AFM cantilever’s spring constant and optical lever sensitivity (OLS) were performed with the Sader method in air ^42^. To calculate the OLS in liquid, a thermal noise spectrum was taken while the cantilever was submerged in imaging media. The super luminescent diode (SLD) was set midway along the cantilever. Using the built-in Igor software (Asylum 15.6.4), the peak was fit and the OLS was determined using the Sader method determined spring constant ^43^.

### Cell culture for vimentin, actin and lysosome 3D imaging

Vimentin−/− HeLa cells stably expressing vimentin-mEmerald were generated as described ^37^. In brief, the endogenous vimentin gene was deleted via CRISPR-Cas9 methods and a verified vimentin−/− clone was stably transduced with a lentiviral vimentin-mEmerald vector. Transduced cells were selected in 500 µg/ml G418 and sorted for uniform mEmerald expression via fluorescence-activated cell sorting. Cells were grown in DMEM-F12 media without phenol red. For experiments, cells were plated on collagen coated 55 kPa, thin (~10 µm) PA gels within 1 cm cloning cylinders (Corning, USA). The cloning cylinder was removed prior to imaging.

### Volume imaging of vimentin, actin and lysosomes

Live HeLa cells expressing vimentin-mEmerald were treated with 1 µM lysotracker dark red (Invitrogen, USA) for 10 min at 37 degrees, washed several times in fresh, equilibrated medium, and then placed on the AFM stage of the optical microscope. Indirect transmitted light was used to find a cell of interest and the prism was positioned next to the cell. The imaging was then switched to side-view. As described in ^23^, a custom MatLab program generated a look-up table for points of best focus. This look-up table was used by custom software in LabView to synchronize the AOTF, camera, mirror and ETLb for volume imaging. Laser power at the sample for live cell imaging was 98 uW for 488 nm and 290 uW for 561 nm. To image F-actin with vimentin, the expressing cells were fixed in 4% formaldehyde for 5 min, stained with alexafluor 568 phalloidin (Invitrogen) for 30 min at 37 deg, washed with PBS, and imaged. Laser power at the sample for fixed cell imaging was 116 uW for 488 nm and 80 uW for 561 nm.

### Generation and use of RAW 264.7 murine macrophage cells expressing F-tractin

RAW 264.7 cells were obtained from ATCC (TIB-71) and maintained in RPMI 1640 medium (ThermoFisher Scientific, cat# 61870036, MA) supplemented with GlutaMAX and 10% heat-inactivated fetal bovine serum (Gemini Bio-Product, CA). Cells expressing F-tractin-Halo were generated using a PiggyBac transposon system. We first produced a stable macrophage cell line expressing reverse tetracycline-controlled transactivator (rtTA) and then used those cells to produce a stable cell line expressing F-tractin-Halo under the control of a tetracycline-dependent promoter. The transfection was performed using the Viromer Red transfection reagent (Lipocalyx, Weinbergweg, Germany) according to the manufacturer’s instructions. Briefly, 2.5 × 10^5^ cells were seeded into one well of a six-well plate, and the cells were transfected the following day. For each well, we used 1.2 µL of Viromer Red transfection reagent, 2.5 µg *piggyBac* transgene plasmid (PB-rtTA or PB-tre—F-tractin—Halo-Hygro) and 0.5 µg of *piggyBa*c transposase plasmid (ratio at 5:1). After 24 hours, the medium was replaced with fresh culture medium and the cells were allowed to recover for 24 hours. Stable transfectants were selected by gradually increasing antibiotic concentrations; geneticin (G418) to a concentration of 500 µg/ml for PB-rtTA and Puromycin to 2.5 µg/ml for PB-F-tractin-Halo. To induce the expression of F-tractin-Halo, the cells were grown in medium containing 50 nM doxycycline for 48 hours. On the day of imaging, cells were plated onto fibronectin coated PA gels within 1 cm cloning cylinders. After one hour, 4 µL of 10^−8^ M JaneliaFluor 549 (JF549) HaloTag ligand was added to the cells for a 30 minute incubation. The dye containing media was replaced with imaging media three times every 10 minutes to removed unbound dye.

### Plasmid construction

pEGFP-C1 F-tractin-EGFP was a gift from Dyche Mullins (Addgene plasmid # 58473; http://n2t.net/addgene:58473; RRID:Addgene_58473). PiggyBac plasmids PB-rtTA and PB-miRE-tre-Puro were kindly provided by Mauro Calabrese (The University of North Carolina at Chapel Hill); PB-rtTA encodes reverse tetracycline-controlled transactivator (rtTA) and G418 resistant gene under UbC promoter ^44^, and PB-miRE-tre-Hygro encodes a protein of interest and a hygromycin resistant gene under tetracycline-dependent and EF1 promoters, respectively. pF-tractin-Halo was first generated by cloning Halo-Tag cDNA (forward primer, aggggggctagcgctcgccaccatggcagaaatcggtactggctttc; reverse primer, cgaagcttgagctcgagatctagtcgactgaattcgcgttatcgc) between AgeI and BglII site of pEGFP-C1 F-tractin-EGFP using Gibson assembly (New England Biolabs, MA). Halo-F-tractin was then amplified using the primers (forward primer, tgaaccgtcagatcgcctggaccggtgccaccatggcgcgaccacgg; reverse primer, aggcacagtcgaaacgcattgtcgacttatggctcgccggaaatctcg), and cloned between AgeI and SalI sites of PB-miRE-tre-Hygro, yielding PB-tre-F-tractin-Halo-Hygro. Those plasmids were confirmed by sequencing before use. The primers were synthesized by Integrated DNA Technologies (CA) and sequences were performed by Genewiz (NJ).

### Fixed light sheet phagocytosis experiment

A 3D printed custom specimen insulating chamber (uploaded as Thingiverse 2035546) was placed on top of the coverslip, with the hole aligned for prism access. The specimen insulator was magnetically secured to the heated microscope stage using Asylum Research’s petri dish clamp. A thin circle of silicone grease (High Vacuum grease, Dow Corning, USA) secured the specimen coverslip to the 3D part as well as functioned to prevent leakage of media. Deionized water was added to a well in the specimen chamber to reduce evaporation. The AFM head with evaporation shield and a calibrated, functionalized cantilever was placed on the scope with the cantilever in the media and allowed to equilibrate.

Using brightfield illumination, a cell was chosen that had lamellipodia and filopodia. The cantilever was lowered to within a few microns above the cell. The prism was brought as close as possible next to the cell and cantilever without touching the cantilever or interfering with the SLD light path. Imaging in side-view then allowed adjustments to cantilever height, light sheet height (using ETLa) and image focus (objective and ETLb). The camera was set to accept external trigger (HCImageLive). Using custom LabView software, the AOTF was synchronized with the camera. The system was triggered by the AFM controller when the force curve began using Asylum Research Software and custom code (IGOR). The sequential timing was 300 ms exposure with a 100 ms delay for both 561 nm and 488 nm lasers. Laser power was chosen to maximize image quality while limiting photobleaching. For 561 nm, this was typically 30 µW of light delivered to the sample, while for 488 nm, this was 15 µW. A virtual deflection curve was taken prior to each experiment. The AFM was set to closed loop Z-position feedback. The trigger distance for the cantilever to travel and stay was chosen so that the bead tip would be within 0.5 µm of the top of the cell. The cantilever was lowered at 1 µm/s. Duration of the experiment was 1800 s dwell toward the cell with a 200 s dwell away from the cell.

### Volume scanning during phagocytosis

Our protocol for initialization of volume scanning is described in our prior report ^23^. Otherwise, the set up for the volume imaging of phagocytosis experiment was similar to the fixed light sheet. The AFM cantilever’s bead tip was positioned over a macrophage and the prism was placed next to the cantilever. After switching to side-view and adjusting the LBS, the volume scan proceeded as described above. AFM settings were the same as in the fixed light sheet experiment. As in the fixed light sheet experiment, the system was triggered to start the experiment with the AFM, synchronizing the force data with the volume imaging.

### Environmental Control

The specimen insulating chamber sits atop the circular sample coverslip. The specimen insulator made use of the Asylum Research petri dish heater and magnetic petri dish clamp while also allowing the prism to freely enter and exit the specimen space. A well that runs along the inside of the specimen insulator above and separate from the specimen media holds DI water to increase humidity of the chamber and discourage evaporation of the media. Additionally, the AFM head is equipped with an evaporation shield that caps the specimen insulating chamber. Once the chamber is enclosed a stable temperature environment (37°C) can be maintained with minimal evaporation.

In addition to the petri dish heater, we used an objective heater (Thorlabs HK-100) with a PIV controller (Thorlabs TC200) to maintain the temperature at the sample location. Settings for both controllers were adjusted until the specimen reached the desired temperature, stability and gradient. The specimen temperature was measured with an Omega thermocouple placed directly in a test media. 37.0°C was achieved with a 0.1°C gradient of the media from edge of the coverslip to the center of the coverslip.

### PA spheres for the PSF in Supplementary Figure 4

PA spheres were generated by the methods as described ^33^, with some minor changes. We used the 3 µm SPG frits to extrude high stiffness PA gel into hexane at 300 RPM stirring. Beads were fixed overnight in AIBN as described ^33^, washed in fresh hexane, dried in air and resuspended into PBS with sonication to separate the beads. The spheres were 12-20 µm in diameter after resuspension, and were coated in a similar manner to the pads above. An aliquot of the PA beads was centrifuged in an Eppendorf tube (7000 RPM, 4 min), and resuspended in 10 mg/mL EDAC in pH 6 MES buffer and rotated for 15 min at room temperature. We centrifuged the beads to remove the EDAC as above, and resuspended them in 10 ug/mL alexafluor 488 goat anti mouse IgG (Invitrogen). After 30 min rotation, the beads were again centrifuged to remove unbound IgG, and resuspended in EDAC (10 mg/mL) with a 10^−6^ dilution of 0.1 µm red fluorescent microspheres (Invitrogen) in PBS. Beads were again rotated for 30 min and then pelleted, and resuspended in plain PBS.

To attach the coated beads to a PA gel pad, low concentrations of beads were suspended onto the gels within the cloning cylinders, and again EDAC was added to 10 mg/ml final concentration. The IgG fluorescence was used to find the PA beads, and then an isolated red fluorescent bead was imaged for the PSF.

### Correlation of Force signal and Local Actin Activity

We performed cross-correlation analysis on our AFM force time series, ***F(t)***, and the local F-tractin fluorescence intensity, ***I(t)***. Regions of interest (ROI) within the fluorescence image stack were selected and the average intensity within the ROI was saved as a time series. The force signal and ROI image intensity signal were then analyzed using cross-correlation code. Both force and intensity data were high pass filtered prior to cross-correlation calculation to remove offset and slope (frequency cut off = 0.5 mHz). For these data, downward forces (in the direction of engulfment) are negative, and upward forces (pushing force of the cell on the bead) are positive. Positive correlations between force and ROI intensity indicate where increased actin and pushing (upward) forces coincide, while negative correlations indicate where higher actin intensity is coincident with downward engulfment forces.

We assumed that the dynamics we observed were not stationary, that is we did not expect the statistics of either of the force or the intensity to be consistent throughout the multistage phagocytic process. Thus we needed to perform our cross-correlation calculations on subsets of the data to assess correlations more locally in time. We selected a time window, *t*_*w*_, over which to make our local assessment of correlation. This window was stepped through the data set at small time increments (1 s) to collect the correlation strength at each experimental time t, as a function of time lag, *τ* (in this case, lag of intensity relative to force).

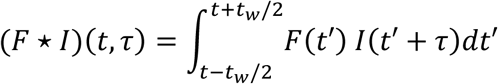

Where (*F* ⋆ *I*) is the unnormalized cross-correlation strength between force and intensity, t is the “zero lag time” over which the correlation is centered, τ is the time lag, and *t*_*w*_ is the time window. We chose not to use a normalized cross-correlation calculation. With normalization (as for Pearson correlation), certain time windows with very small but correlated intensity fluctuations yielded cross-correlation strength comparable to those time windows with very large signal fluctuation correlation, falsely emphasizing the smaller signals’ significance. With no normalization, correlation strength was proportional to signal magnitude. This emphasized correlations between larger, and in our view, more relevant fluctuations in force and actin intensity.

## ACKNOWLEGEMENTS

R.S. acknowledges funding from NIH and NSF (NSF/NIGMS 1361375).

R.S and K.M.H acknowledge NIH funding (NIBIB P41-EB002025).

M.B. would like to recognize NIH grants 5R01GM118847-03 and 1R01NS111588-01A1 for providing funding.

E.N. and M.E.K. were supported on NIH 5T32GM008570-19.

C.M.H. was supported by the NSF GRFP (DGE-1650116)

## AUTHOR CONTRIBUTIONS

E.N. performed all AFM-LS experiments and analysis not noted otherwise with assists from M.E.K and C.M.H. C.M.H. performed all two-color volumetric imaging on HeLa cells and related analysis. M.R.F. performed correlation analysis.

M.B and B.M.C performed transfections on HeLa cells and helped design vimentin experiments, with assists from E.T.O on cell culture.

E.T.O with help from Jacob Brooks fabricated the PA beads for the PSF testing. E.N. developed the PSF theory and C.M.H collected the PSF data. E.N. performed the AFM noise testing and spectral analysis with assists from M.R.F.

R.S. conceived and designed the AFM-LS system.

R.S. and M.R.F. conceived and designed biophysics experiments using the AFM-LS system. S.G. helped in development of biophysical hypotheses and experimental design for phagocytosis experiments.

E.N. and C.M.H. built and tested the AFM-LS system.

E.T.O. prepared gel substrates and handled most cell culture activities.

J.H. wrote and implemented all control code for the AFM-LS system with assists from C.M.H. and E.N.

M.E.K., T.W. and K.M.H. designed and performed all plasmid construction and genetic modification of RAW 264.7 cells, with assists from E.T.O.

All authors wrote the manuscript.

## SUPPLEMENTARY VIDEO CAPTIONS

**Supplementary Video 1: Fast volumetric time-lapse of lysosomes labeled with Lysotracker Deep Red (magenta) and vimentin-mEmerald (green) in a live HeLa cell.** 75 slices per color, 5 ms exposure time, 5 ms readout time, 100 ms between volumes, 1.5 s per volume. Voxel size: 106 × 106 × 220 nm. Total volume: 29.3 × 26.3 × 16.5 µm.

**Supplementary Video 2: Volumetric image of a fixed HeLa cell with labeled actin-Phalloidin (magenta) and vimentin-mEmerald (green).** 300 slices per color, 5 ms exposure time, 5 ms readout time, 6 s per volume. Voxel size: 106 × 106 × 108 nm. Total volume: 52.6 × 19.1 × 32.4 µm.

**Supplementary Videos 3 and 4: AFM Force curve and side view time-lapse of a RAW 264.7 cell stably expressing Halo-F-tractin labeled with JF 549 during phagocytosis.** *(Top)* Beaded AFM cantilever tip labeled with Goat pAb to Mouse IgG AlexaFluor488 (channel not shown). 300 ms exposure with 100ms delay between channels. Every 10^th^ frame is shown. Scale bar: 5 µm. (*Bottom*) The vertical, red line indicates the force data for the corresponding image frame. Imaging includes the dwell and the retraction of the cantilever, though the force data associated with the retraction of the AFM is not shown due to large adhesive forces drastically changing the scale.

**Supplementary Video 5: Volumetric time-lapse of a RAW 264.7 cell stably expressing Halo-F-tractin labeled with JF 549 during phagocytosis.** *(Top)* 75 slices per color, 10 ms exposure time, 5 ms readout time, 2.85 s per volume, 7 s between volumes. Voxel size: 106 × 106 × 200 nm, Total volume: 28.1 × 21.7 × 19.0 µm. (*Bottom*) The vertical, red line indicates the force data for the corresponding image frame.

**Supplementary Video 6: Podosome-like structures visible in the maximum intensity XY projection of the phagocytic cup from Figure 4**.

