## Supplementary Figures for "Combined Atomic Force Microscope and Volumetric Light Sheet System for Mechanobiology"

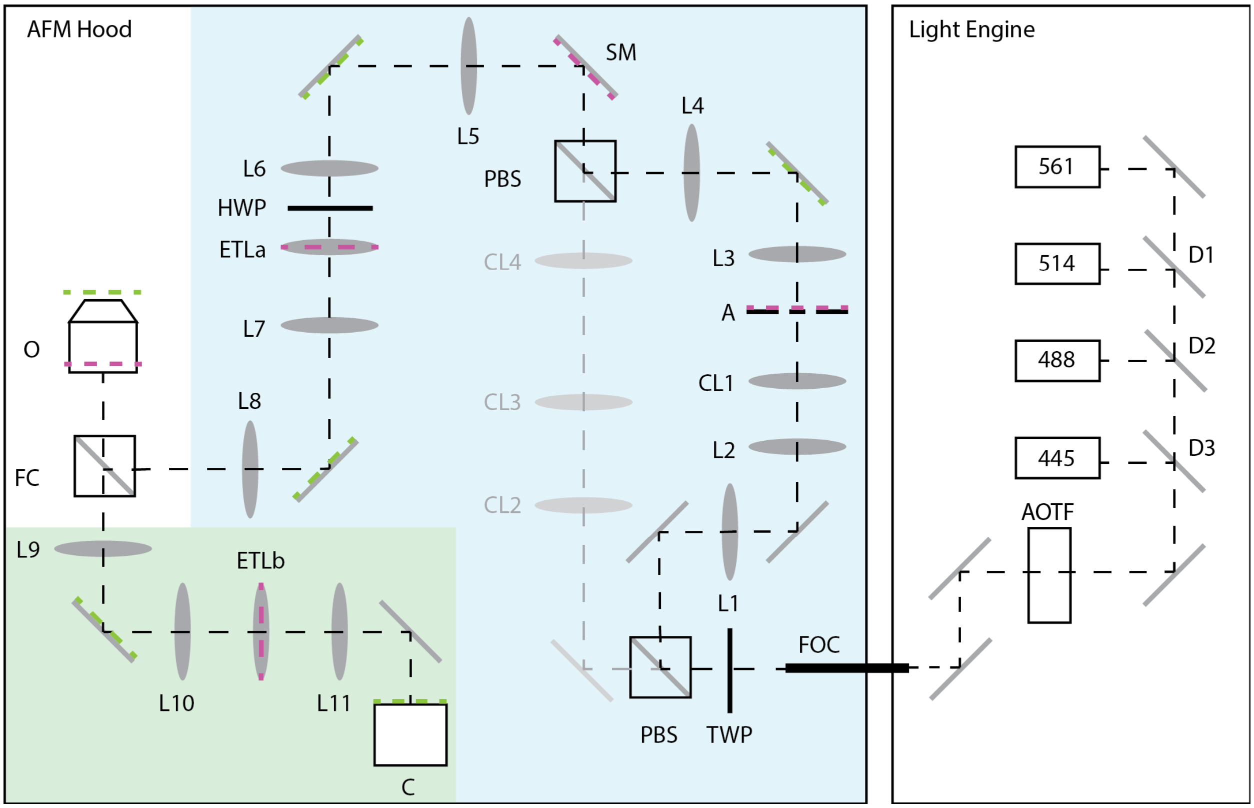


**Supplementary Figure 1 | Optical System Schematic.** L1-L11: Lenses; CL1-CL4: Cylindrical Lenses; D1-D3: Dichroics; FOC: Fiber Optic Cable; TWP: Tunable Wave Plate; PBS: Polarizing Beam Splitter; A: Annulus (Inner Radius = 0.1 mm, Outer Radius = 0.5 mm); SM: Steering Mirror; HWP: Half Wave Plate; ETLa-ETLb: Electrically Tunable Lenses; FC: Filter Cube; O: Objective Lens (UplanSAPO 60x/1.2 NA W, Olympus); C: Camera (Hammamatsu Orca Flash 4.0 v2). Focal Length of Lenses: (L1) 25 mm, (L2) 150 mm, (L3) 100 mm, (L4) 125 mm, (L5) 125 mm, (L6) 125 mm, (L7) 125 mm, (L8) 175 mm, (L9) 175 mm, (L10) 125 mm, (L11) 125 mm, (CL1) 50 mm, (CL2) 14.5 mm, (CL3) 150 mm, (CL4) 100 mm. Green and Magenta dashed lines represent sample and pupil conjugate planes respectively.

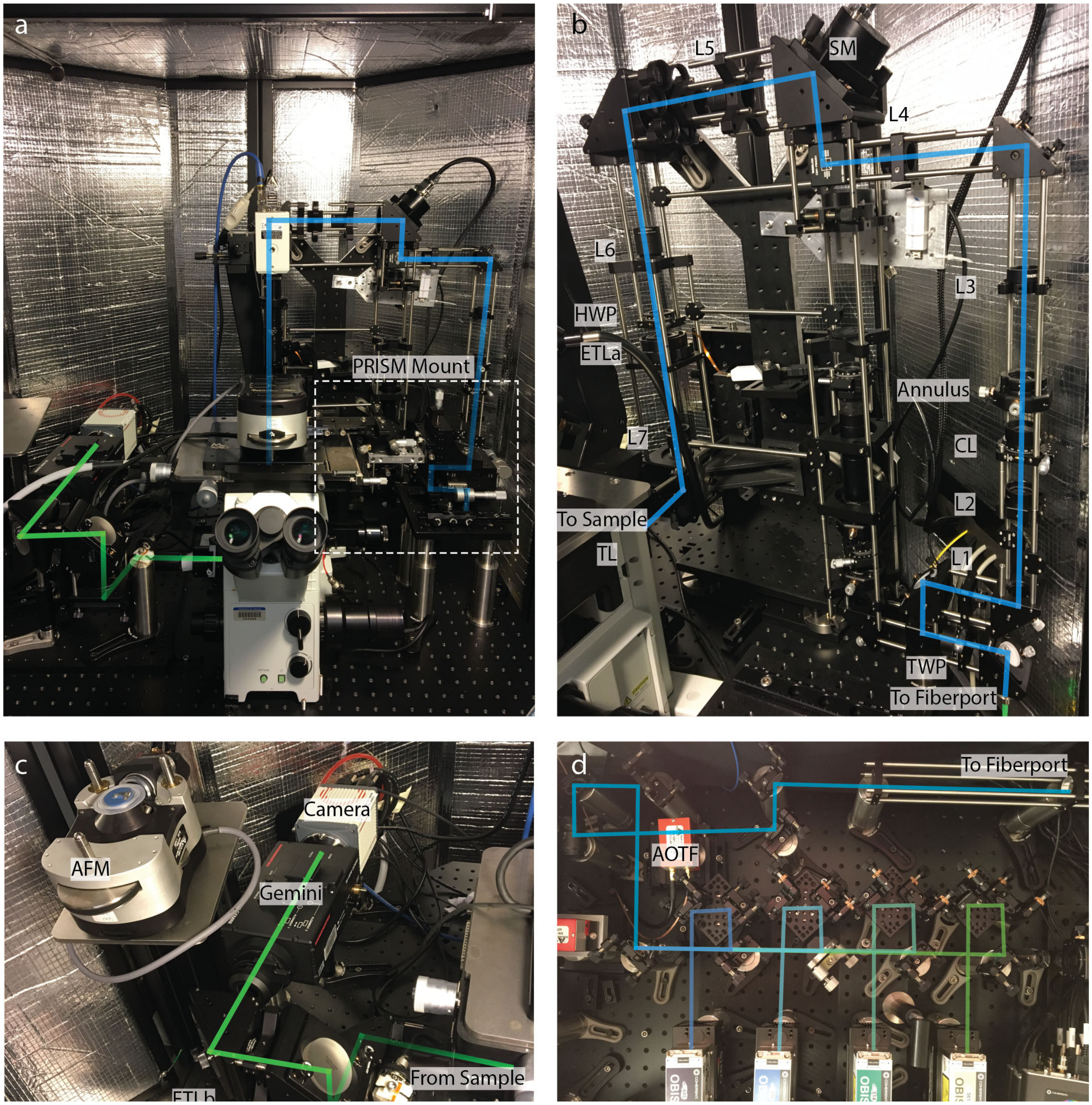


**Supplementary Figure 2 | AFM-LS System.** Our system has been designed to enclose the beam shaping optics, the Olympus confocal microscope, and the AFM into a mechanical noise isolation chamber situated on top of a vibration isolation stage. **a**, The interior of the chamber showing the up-right excitation path traced in cyan, the Olympus inverted microscope, the emission path traced in green, the AFM positioned onto the microscope stage, and the PRISM mount. **b**, A detailed view of the excitation path. The laser light is shaped into a line Bessel sheet after passing through the cylindrical lens (CL) and annulus. The position of the light sheet is controlled by the fast steering mirror (SM) and the electrically tunable lens (ETLa) which are conjugate to the back focal plane of the objective. See Supplement Figure 1. **c**, Zoomed-in view of the emission path to the camera. The AFM head is inverted onto a shelf added to the chamber. A fiber optic cable connects the optics in this isolation chamber to a separate optics chamber housing the lasers and AOTF. **d**, The light engine: A photo showing the top-down view of the four lasers directed into an AOTF and then coupled into a fiber optic.

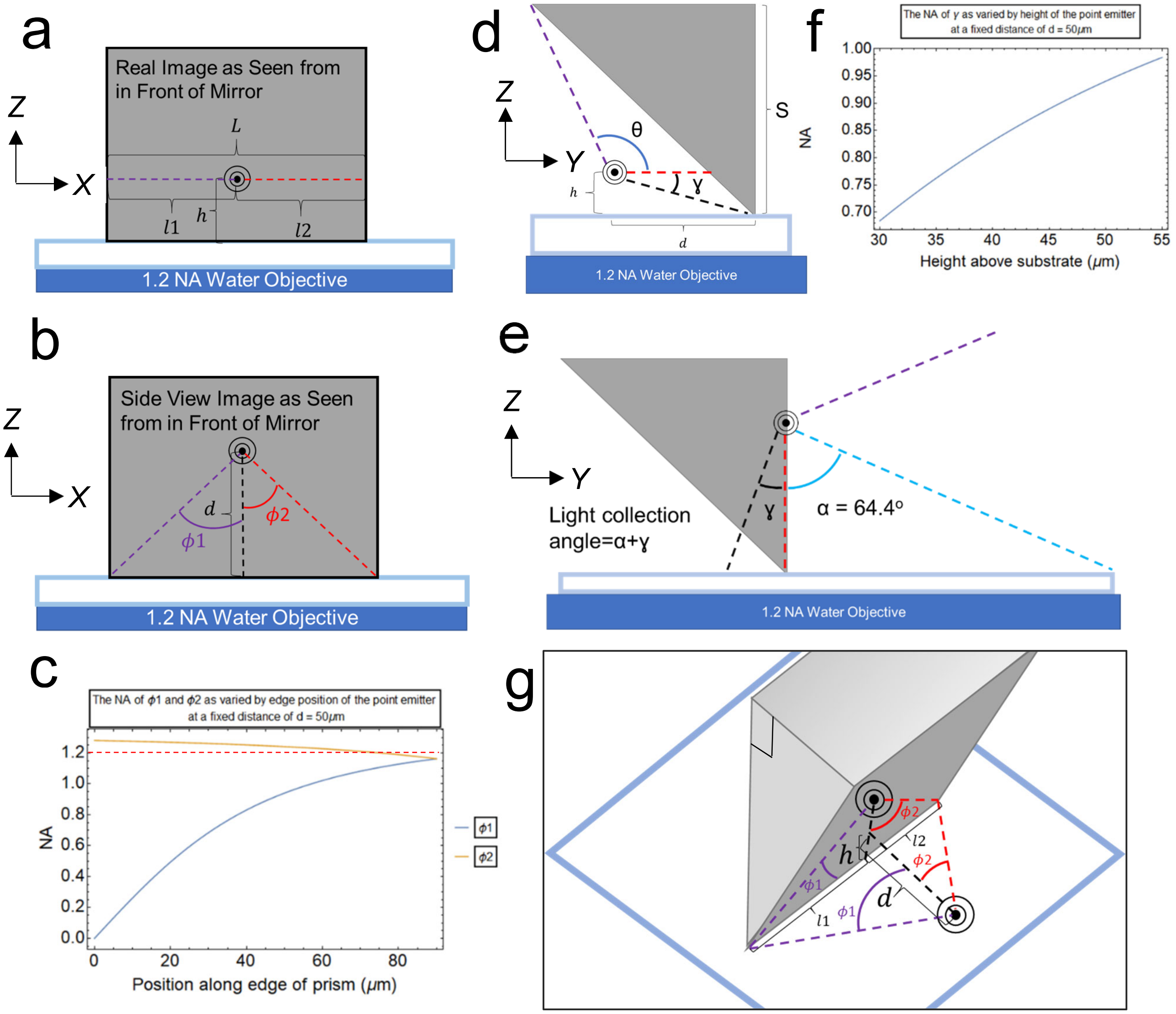


**Supplementary Figure 3 | Light collection of the prism**. **a,** A diagram showing the x-z imaging plane of a point emitter at a distance, $d,$ away from the mirror with partitions $l1$ and $l2$. The height, $h,$ above the substrate is shown for orientation. **b,** The point emitter is rotated 90° when imaged in side view, forming the two angles of the partitions, $\phi1$ and $\phi2$. The height above the substrate of the virtual image is the distance, *d*, from the prism of the point emitter. **c,** Numerical aperture (NA) plot for $\phi1$ and $\phi2$ at fixed distance 50μm as the point emitter is varied along the 180 μm prism edge. The red dashed line is the NA of the objective lens, above which light collection does not improve. ($\phi1 = acrtan\left(\frac{l1}{d}\right)$; $\phi2 = acrtan\left(\frac{l2}{d}\right)$; NA = 1.33 sin(angle)). NA in the x-axis is the mean of NA $\phi1$ and NA $\phi2$ **d,** A diagram showing the y-z imaging plane of a point emitter a distance, *d,* away from the mirror and height, *h,* above the substrate. **e,** The point emitter is rotated 90° when imaged in side view. θ becomes limited by the angle that the objective can collect, in this case α = 64.4°, while ɣ is typically less than α. **f,** Numerical aperture plot for $\gamma$ at fixed distance 50 μm (ɣ = $arctan(\frac{h}{d})$; NA = 1.33 sin(angle)). The other angle is objective lens limited (NA=1.2). NA in the z-axis is the mean of these two NAs. **g,** A three-dimensional graphic showing dimensions of the point emitter for the x-z plane.

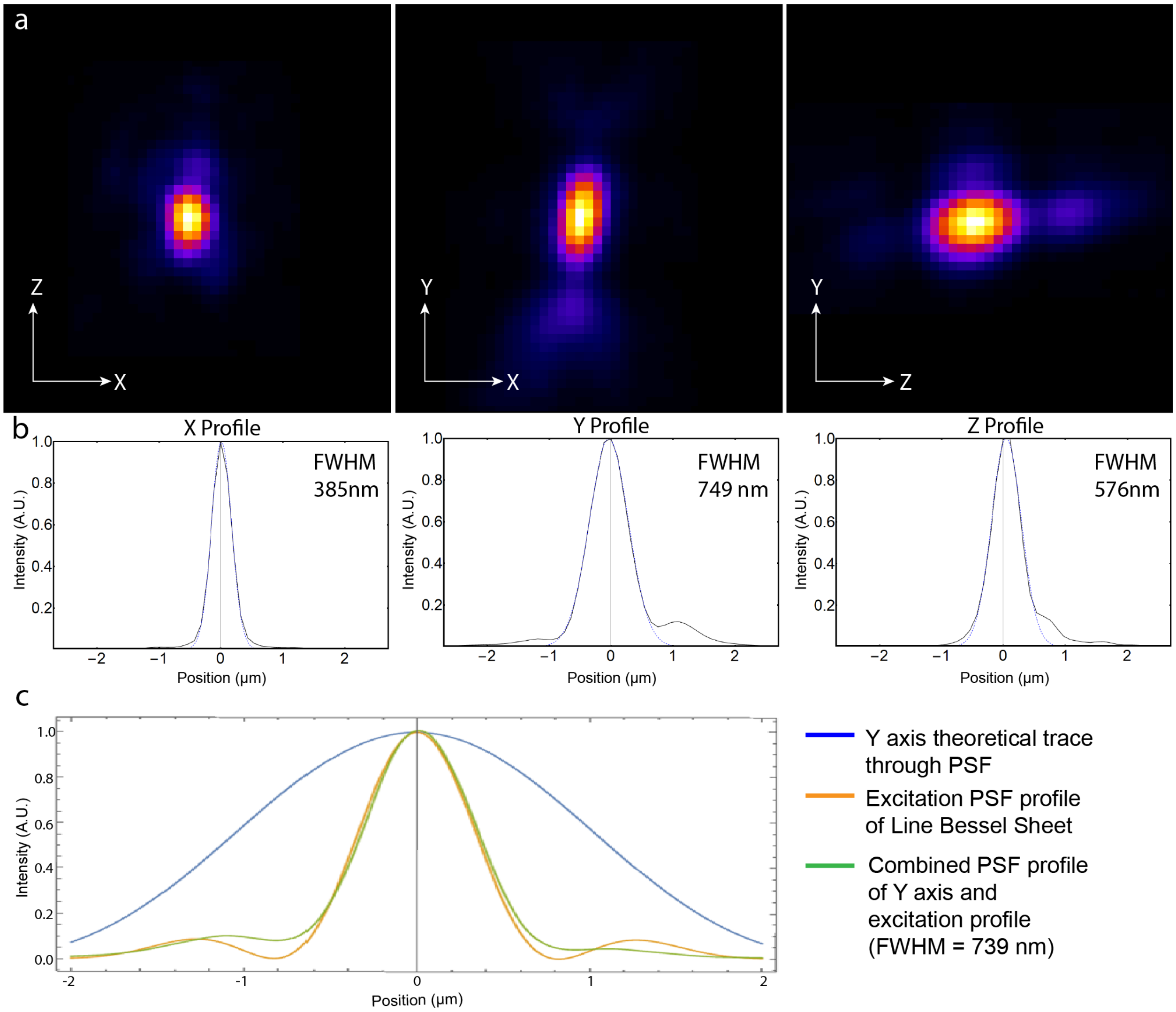


**Supplementary Figure 4 | Point spread function of side view imaging. a,** Optical sections of the PSF in the x-y plane, the x-z (side view) plane, and the y-z plane. **b,** The plotted profiles of each axes through the center of the PSF. **c,** The theoretical calculated plot for the Y direction. The blue dotted lines in the profile plots are the gaussian fit to each plot from which the FWHM is determined. X is the direction parallel to the prism edge, Z is the direction from substrate to top of cell and Y is the light sheet stepping direction. Using the theory presented above (Supplementary Figure 3) and the Rayleigh criterion for resolution, we calculate a x-axis resolution of 381 nm and a z-axis resolution of 574 nm with values based on the bead position with respect to the prism and substrate (d = 77 μm, h = 5 μm, $l1$ = 60 μm, $l2$ = 120 μm, ʎ = 605 nm). The expected axial (y-axis) PSF profile is calculated using these values then combined with the PSF profile of the Line Bessel sheet (Combined FWHM= 739 nm). Experimentally determined values are in excellent agreement with theory. The distance where the PSF was taken represents the expected poorest imaging quality for experimental conditions. Closer to the prism (d = 10 μm), x resolution is objective lens limited (res ~ 310 nm), and z resolution is ~ 400 nm. Typical cell distance to the prism with the AFM in place is between 50 μm and 70 μm. See Online Methods for creation of PSF sample.

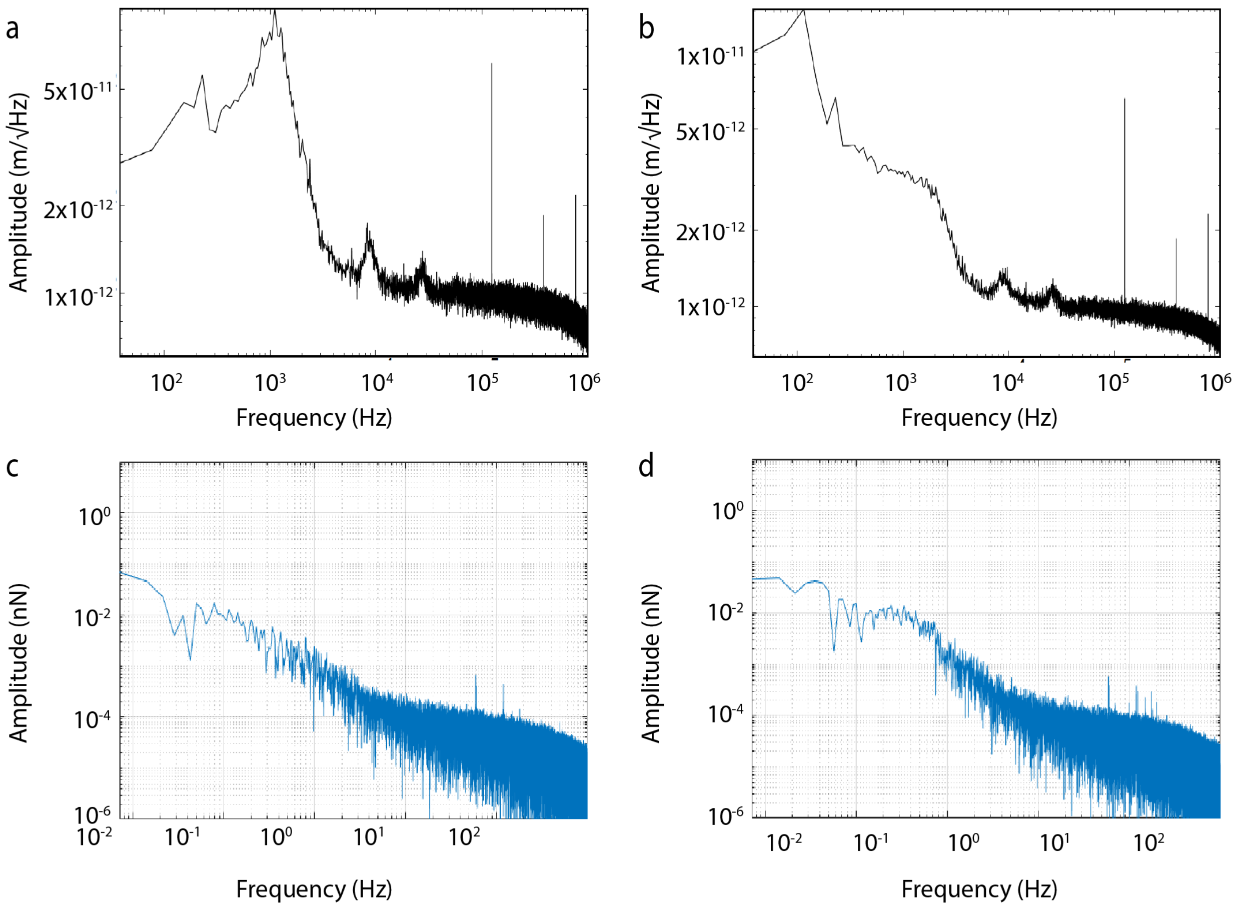


**Supplementary Figure 5 | Thermal frequency spectra and noise determination in AFM. a**, Thermal noise spectrum collected over a 1 MHz range of frequencies for a cantilever near the gel substrate at 37°C. The main peak around $10^3$ Hz is the thermal contribution from the cantilever. **b,** Thermal noise spectrum collected over a 1 MHz range of frequencies for a cantilever in contact with the surface and with volume scanning at 37°C. For collection at 2 kHz, the bandwidth at which experiments are collected, the total $x_{RMS}$ noise is 0.189 nm which corresponds to a noise level of ~16 pN. **c**, Fourier Transform of fixed z-position time series without volume scanning optics running. This shows the Fourier Transform of a fixed z-position AFM time series of a cantilever engaged with a gel substrate at 37°C. **c,** Fourier Transform of fixed z-position time series with volume scanning optics running. This shows the same cantilever in **c** engaged with the gel substrate and with volume scanning. Data collection for both **c** and **d** was taken at 2 kHz bandwidth for 65 s. For optical system parameters used for volumetric imaging, we expected noise at frequencies at 0.4 Hz, 25 Hz and 200 Hz. Contributions to the noise at 0.4 Hz and 200 Hz are seen in the right hand curve. However, they are on the order of single pN or less, which is an order of magnitude lower than the theoretical thermal noise floor of the cantilever. The two peaks that are apparent at high frequencies in both no scanning **c** and scanning **d** occur at 60 Hz and 120 Hz, indicating some small electronic noise is present.
